# Graded Notch Signaling Functions as a Rheostat of Lineage Plasticity and Therapy Resistance in Prostate Cancer

**DOI:** 10.1101/2025.11.23.690056

**Authors:** Yuyin Jiang, Siyuan Cheng, Catherine Yijia Zhang, Xiao Jin, Longjun Li, Melanie Fraidenburg, Choushi Wang, Isaac Yi Kim, Ping Mu

## Abstract

Resistance to androgen receptor (AR)-targeted therapies such as enzalutamide in castration-resistant prostate cancer (CRPC) often arises through lineage plasticity, yet the molecular mechanisms that define this process remain incompletely understood. While previous studies reported that Notch1 and Notch2 exert distinct and sometimes opposing effects in prostate cancer differentiation, the integrated role of Notch pathway activity has not been systematically explored. Here, we identify Notch signaling as a graded Rheostat that governs prostate cancer cell fate transitions. Integrative transcriptomic and functional analyses revealed that intermediate Notch activity maintains a stem-like progenitor state, whereas reduced or elevated signaling drives divergent differentiation trajectories toward luminal or neuroendocrine lineages, respectively. During CRPC progression and enzalutamide resistance, Notch signaling becomes dynamically rewired, peaking in progenitor-like populations that sustain plasticity and survival. Both CRISPR-mediated knockout and pharmacologic inhibition of Notch signaling depleted these progenitor cells and restored enzalutamide sensitivity. These findings demonstrate that the level, rather than the binary presence, of Notch signaling dictates lineage directionality and therapeutic response in CRPC, establishing it as a tunable and actionable driver of resistance.

## Introduction

Prostate cancer remains a leading malignancy in men and is primarily driven by androgen receptor (AR) signaling, making this pathway a central therapeutic target[1]. Androgen deprivation therapy (ADT), which reduces androgen levels or inhibits AR activity, has long been the cornerstone of treatment for advanced disease. Although ADT elicits strong initial responses, resistance inevitably develops, leading to castration-resistant prostate cancer (CRPC)—a more aggressive state that progresses despite androgen depletion [1]. The challenge of CRPC spurred the development of next-generation AR-targeted therapies, such as enzalutamide and apalutamide, which provide more potent and comprehensive AR signaling inhibition and became the standard-of-care for this patient population [2, 3]. However, the clinical benefits of these advanced therapies are also frequently curtailed by the development of acquired resistance [4-6]. Addressing this resistance to advanced AR-targeted agents in CRPC is therefore a critical focus of ongoing oncological research and underscores the urgent need for novel therapeutic strategies.

Several molecular mechanisms contribute to resistance to AR-targeted therapies, including restoration of AR-driven transcriptional programs and activation of alternative signaling pathways that bypass AR dependency [7]. Emerging evidence has also identified lineage plasticity, in which prostate cancer cells escape the luminal epithelial lineage and adopt a multi-lineage, progenitor-like state independent of AR signaling [8-11]. Lineage plasticity has emerged as a central adaptive process underlying therapy resistance across diverse cancers, including prostate, breast, lung, pancreatic cancers, and melanoma [9, 11-17], and characterized by a spectrum of genomic and transcriptional aberrations [12, 14, 18-33]. For instance, our previous work showed that concurrent loss of TP53 and RB1 promotes treatment resistance in prostate cancer partly through activation of JAK–STAT signaling [9, 11]. Notably, many resistance mechanisms involve cancer cells acquiring stem-like features, suggesting that stemness-associated pathways may directly drive lineage plasticity. However, how prostate cancer cells sustain this intermediate stem-like state, maintaining plasticity without fully transdifferentiating into alternative lineages such as neuroendocrine prostate cancer (NEPC), remains poorly understood.

Among these, Notch is a central stemness-associated signaling pathway. It is a highly conserved cell– cell communication system that regulates key processes such as cell fate determination, proliferation, differentiation, and apoptosis, essential for both development and tissue homeostasis [34, 35]. In human cells, ligands of the Delta-like (DLL1, DLL3, DLL4) and Jagged (JAG1, JAG2) families engage one of the four Notch receptors (NOTCH1–4) on adjacent cells, triggering proteolytic cleavages that release the Notch intracellular domain. This domain translocates to the nucleus and forms a transcriptional complex with CSL and co-activators to regulate target genes including *HES* and *HEY* (Figure 1). Owing to its broad regulatory functions, aberrant Notch signaling has been widely implicated in the pathogenesis of human cancers [36]. In prostate adenocarcinoma, Notch signaling has been primarily characterized as oncogenic, with pathway activation promoting cell proliferation, survival, migration, and invasion [37]. However, this view has been complicated by emerging evidence that Notch can exert *opposing* effects depending on cellular context. During neuroendocrine transdifferentiation from adenocarcinoma, Notch signaling undergoes a striking functional reversal, from an oncogenic driver to a tumor-suppressive barrier that constrains neuroendocrine differentiation [38].

**Figure 1.**
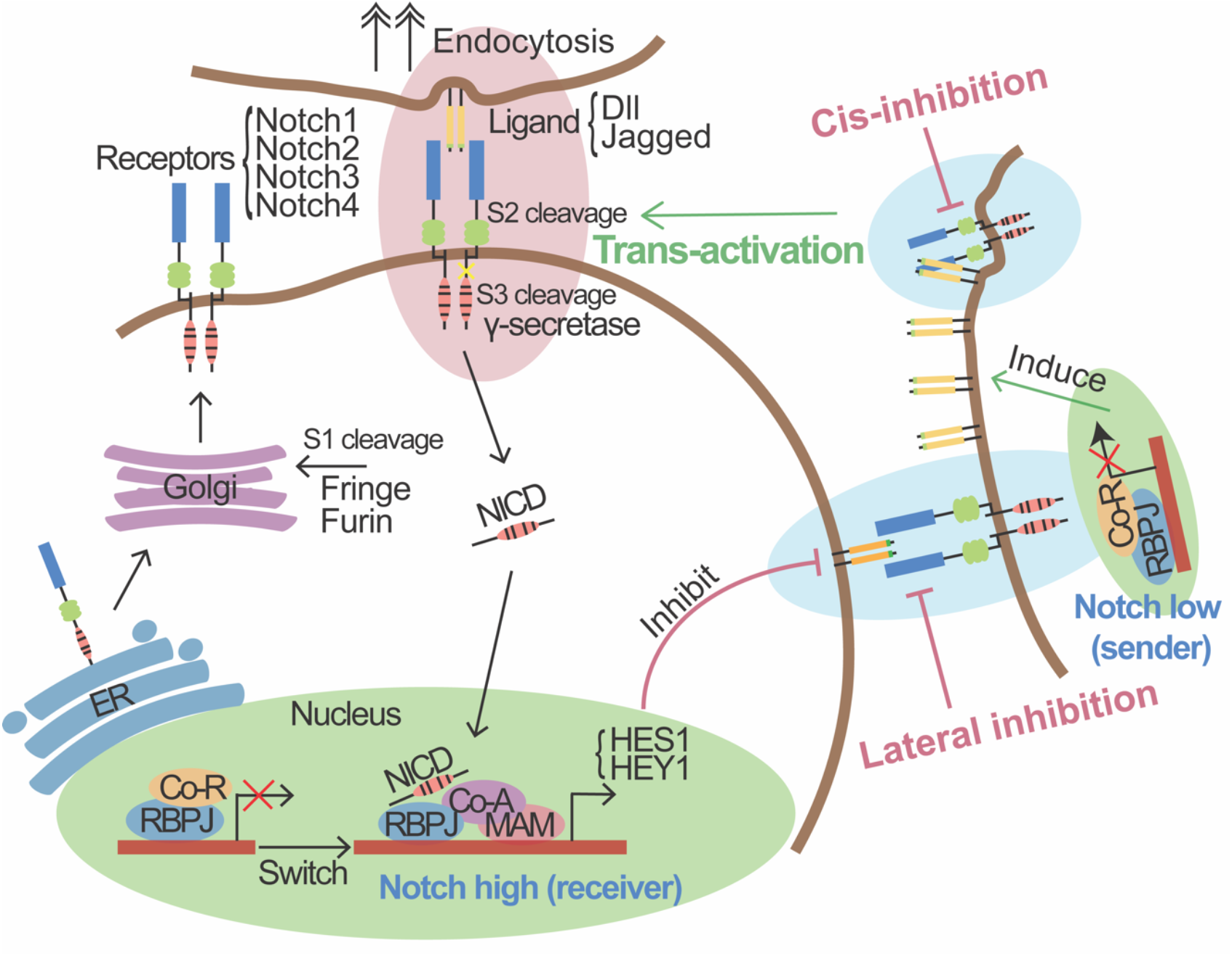
Complex and Context-Dependent Roles of Notch Signaling. Notch receptors, after S1 cleavage, form heterodimer and traffic to the cell surface. Trans-activation occurs when ligands on an adjacent cell bind a receptor, inducing S2 and S3 cleavages. This releases the NICD, which translocates to the nucleus and converts RBPJ from a transcriptional repressor into an activator by recruiting co-activator (Co-A) proteins like MAM, stimulating target genes such as *HES1* and *HEY1*. This can induce lateral inhibition in neighboring Notch^low^/Ligand^high^ sender cells. Cis-inhibition occurs when ligands and receptors on the same cell interact, attenuating signaling. Co-R: Co-repressor; ER: Endoplasmic Reticulum; MAM: Mastermind-like.

In this study, we uncover a graded, rheostat-like role of Notch signaling in regulating lineage plasticity and therapeutic resistance in prostate cancer. We demonstrate that TP53/RB1 co-deficiency activates Notch signaling, positioning Notch as a downstream effector that maintains a stem-like progenitor state and sustains enzalutamide resistance. Mechanistically, intermediate Notch activity preserves a progenitor-like equilibrium, whereas reduced or excessive signaling drives divergent differentiation toward luminal or neuroendocrine lineages, respectively. Genetic or pharmacologic inhibition of Notch disrupts this balance, promoting lineage reprogramming and restoring drug sensitivity. Collectively, these findings identify Notch signaling as a dynamic and tunable mediator linking TP53/RB1 loss to lineage plasticity and therapy resistance, highlighting it as a promising therapeutic target to overcome resistance in advanced prostate cancer.

## Results

### Dynamic Rewiring of Notch Signaling Across Prostate Cancer Progression

Notch signaling exhibits inherent heterogeneity across cell populations due to mechanisms such as trans-activation, cis-inhibition, and lateral inhibition (Figure 1). Hence, we started our characterization of Notch signaling at single-cell transcriptomic level including four distinct cell populations: castration-sensitive prostate cancer adenocarcinoma (CSPC^AD^), castration-resistant prostate cancer adenocarcinoma (CRPC^AD^), CRPC progenitor (CRPC^Progenitor^), and CRPC neuroendocrine (CRPC^NE^) (Figure 2A). Visualization of the calculated Notch pathway activity score onto this UMAP revealed a heterogeneous distribution of Notch signaling across these populations, with notably higher activity in majority of the CRPC^AD^ and CRPC^progenitor^ compared to the most naive and early stage treatment-sensitive prostate cancer (CSPC) clusters (Figure 2B). Quantification confirmed the gradual elevation of Notch signaling activity from CSPC to CRPC^AD^ and then to CRPC^Progenitor^. Consistent with previous report, the Notch signaling dropped in the trans-differentiated neuroendocrine prostate cancer cells (Figure 2C). To further determine whether this elevated Notch activity observed during the progression to castration-resistant prostate cancer reflects an intrinsic adaptive response of the luminal epithelial cells-of-origin to androgen deprivation, we then examined Notch signaling dynamics in the wild-type mouse luminal epithelial cells throughout the castration and regeneration cycle [38]. We found that mouse luminal epithelial cells exhibited a progressive increase throughout the castration phase, a period directly analogous to androgen deprivation therapy. Following the reintroduction of androgens, this elevated Notch activity then gradually declined as the prostate underwent regeneration (R2D-R28D) (Figure 2D). To further validate these observations from single-cell analyses with a larger sample number, we analyzed bulk RNA-seq data from human patient samples. This broader dataset corroborated the trend of differential Notch activity observed in the single-cell studies that increase along with cancer progression and resistance but decreased in neuroendocrine prostate cancer cells (Figure 2E).

**Figure 2.**
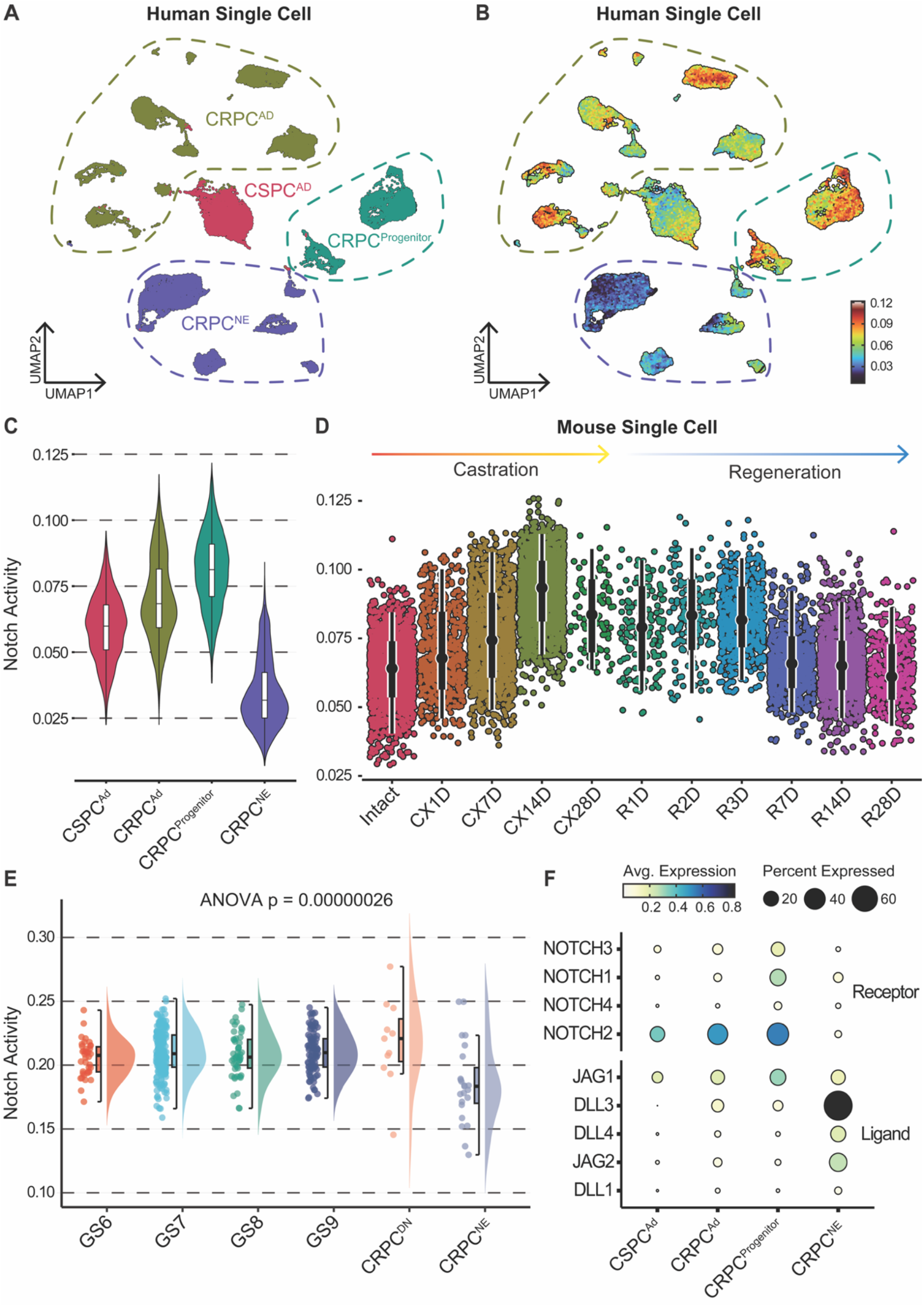
Dynamic Rewiring of Notch Signaling During Prostate Cancer Progression. (A) UMAP plot of human single-cell RNA sequencing data, illustrating distinct prostate cancer cell populations: castration-sensitive prostate cancer adenocarcinoma (CSPC^AD^), castration-resistant prostate cancer adenocarcinoma (CRPC^AD^), CRPC progenitor (CRPC^Progenitor^), and CRPC neuroendocrine (CRPC^NE^). (B) UMAP plot from (A) colored by Notch activity score, with warmer colors (red/orange) indicating higher activity and cooler colors (blue) indicating lower activity. (C) Violin plots comparing Notch pathway activity scores across the indicated human prostate cancer cell populations. (D) Violin and scatter plots depicting Notch pathway activity in mouse single-cell RNA-seq dataset over a time course of castration (CX1D to CX28D) and subsequent regeneration (R1D to R28D), compared to intact controls. (E) Violin plots showing Notch activity across prostate cancer patient groups. GS: Gleason score; CRPC^DN^: CRPC: AR^-^/NE^-^. (F) Dot plot illustrating the average expression and percentage of cells expressing key Notch receptors and ligands within the distinct human prostate cancer cell populations.

Having established and validated these dynamic changes in overall Notch pathway activity across different disease states and model systems, our next step was to identify the specific Notch signaling components (e.g., ligands, receptors, and target genes) responsible for these alterations by interrogating their expression at the single-cell level within our defined prostate cancer populations. The dot plot (Figure 2F) revealed a distinct and consistent trend in the expression changes of Notch receptors and ligands, aligning with the calculated Notch signaling activity. Three out of four Notch receptors were detected in prostate cancer cells, with their expression increasing progressively from CRPC^AD^ to CRPC^Progenitor^, and then decreasing in CRPC^NE^, as expected. Interestingly, the expression of Notch ligands, including members of the JAG and DLL gene families, exhibited a divergent pattern and were notably upregulated in CRPC^NE^, where Notch receptor expression was lower. This seemingly paradoxical observation is consistent with intrinsic Notch regulatory feedback mechanisms, such as lateral inhibition, wherein active Notch signaling often suppresses ligand expression, confirming the previous characterized Notch activity changes.

### Notch Activity Peaks in Progenitor-like States and Correlates with Lineage Plasticity

To further characterize Notch activity, we first compared the expression of Notch pathway components and lineage markers across the parental LNCaP/AR prostate cancer cell line and three distinct lineage-plastic cell models: the enzalutamide-resistant gTP53/RB1 double knockout model (gTP53/RB1-R), LASCPC1, and PC3 (Figure 3). As shown in Figure 3A, the lineage-plastic cell lines, particularly the stem-like gTP53/RB1-R and PC3 models, show significantly elevated expression of Notch pathway components, including *NOTCH2* and the downstream effectors *HES1* and *HEY1*, compared to the LNCaP/AR cells. In contrast, the neuroendocrine-like LASCPC1 cell line showed decreased *NOTCH2* expression. This result confirmed the trend observed in the human single-cell RNA sequencing data, where Notch activity is elevated in progenitor CRPC and suppressed in neuroendocrine CRPC. Furthermore, analysis of lineage marker expression confirmed that the gTP53/RB1-R, LASCPC1, and PC3 models exhibit significantly higher expression of markers associated with stem-like (*SOX2, KRT7*), Epithelial-to-Mesenchymal Transition (EMT) (*CDH2, EPAS1*), and Neuroendocrine-like (NE-like) states (*ASCL1, SYP, NSE, CHGA*), confirming their lineage-plastic phenotype. Conversely, AR signaling (*TMPRSS2, HERC3*) and Luminal markers (*CLDN3, PSCA*) were generally suppressed in these models (Figure 3B). This data supports a strong correlation between elevated Notch signaling and the stem-like/progenitor state of lineage-plastic, enzalutamide-resistant prostate cancer. We corroborated the expression of key Notch components across a broader panel of human prostate cancer cell lines using bulk RNA-sequencing data (Figure 4A-D). Western blot analysis further validated the protein expression of NOTCH2, DLL3, and HES1 in selected lineage-plastic cell lines (PC3, gTP53/RB1-R, and LASCPC1), confirming the gene expression trends (Figure 4E).

**Figure 3.**
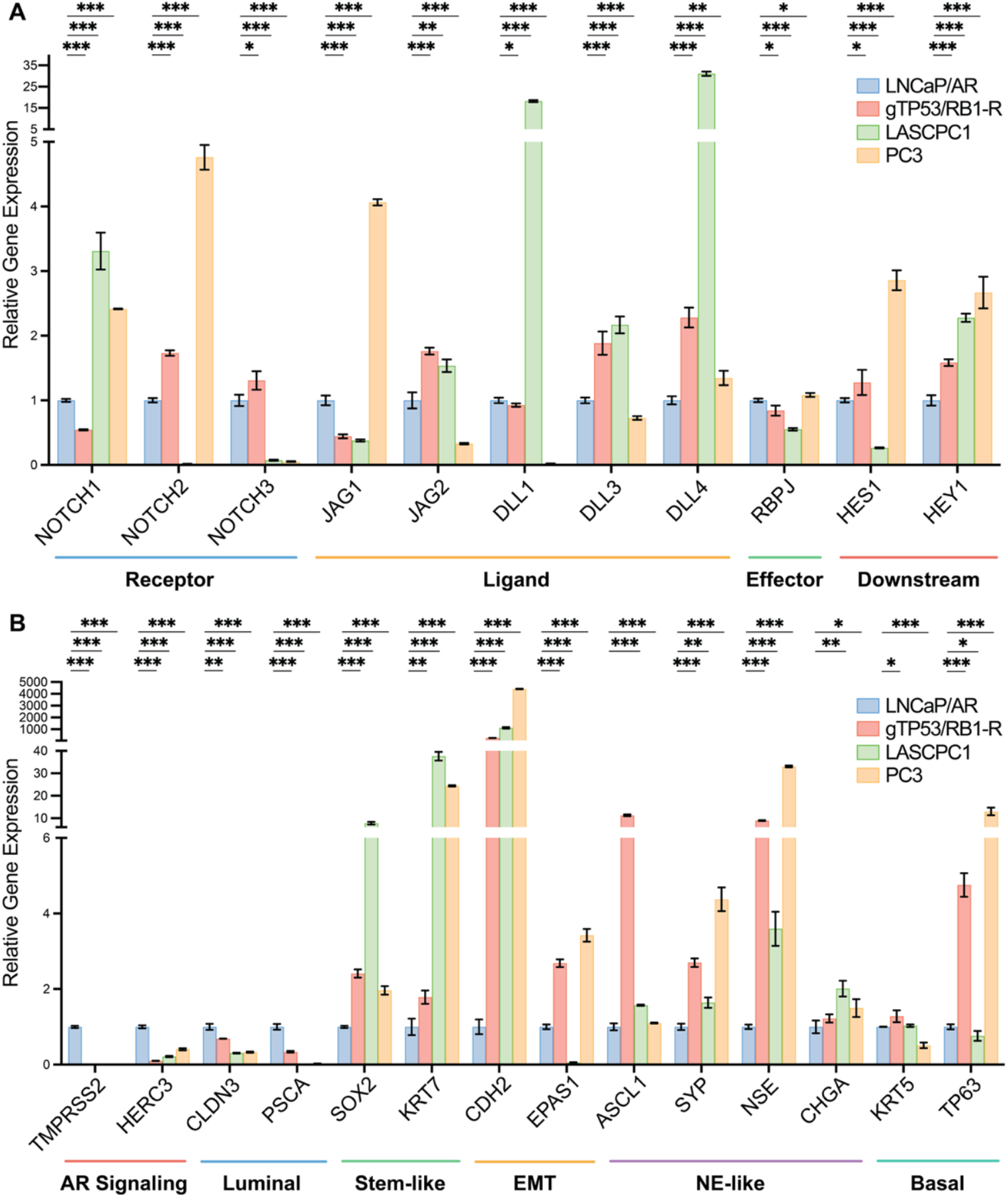
Notch Signaling Peaks in Lineage-Plastic, Stem-like States of Prostate Cancer Cells. (A) Relative gene expression of key components in the Notch signaling pathway (Receptors, Ligands, Effector, and Downstream genes) across the parental LNCaP/AR cell line and three distinct lineage-plastic cell models: the enzalutamide-resistant gTP53/RB1 double knockout model (gTP53/RB1-R), LASCPC1, and PC3. The stem-like gTP53/RB1-R and PC3 cell lines show higher expression of *NOTCH2* as well as the downstream effectors *HES1* and *HEY1* while the Neuroendocrine-like LASCPC1 cell line shows decreased *NOTCH2* expression. (B) Lineage marker gene expression profiles in the same cell lines. The gTP53/RB1-R, LASCPC1, and PC3 models exhibit significantly higher expression of markers associated with stem-like (*SOX2, KRT7*), Epithelial-to-Mesenchymal Transition (EMT) (*CDH2, EPAS1*), and Neuroendocrine-like (NE-like) (*ASCL1, SYP, NSE, CHGA*) states, confirming their lineage-plastic phenotype. Conversely, AR signaling (*TMPRSS2, HERC3*) and Luminal markers (*CLDN3, PSCA*) are generally suppressed. p-values are calculated by multiple t-tests. Significance is denoted as *p < 0.05, **p < 0.01, ***p < 0.001.

**Figure 4.**
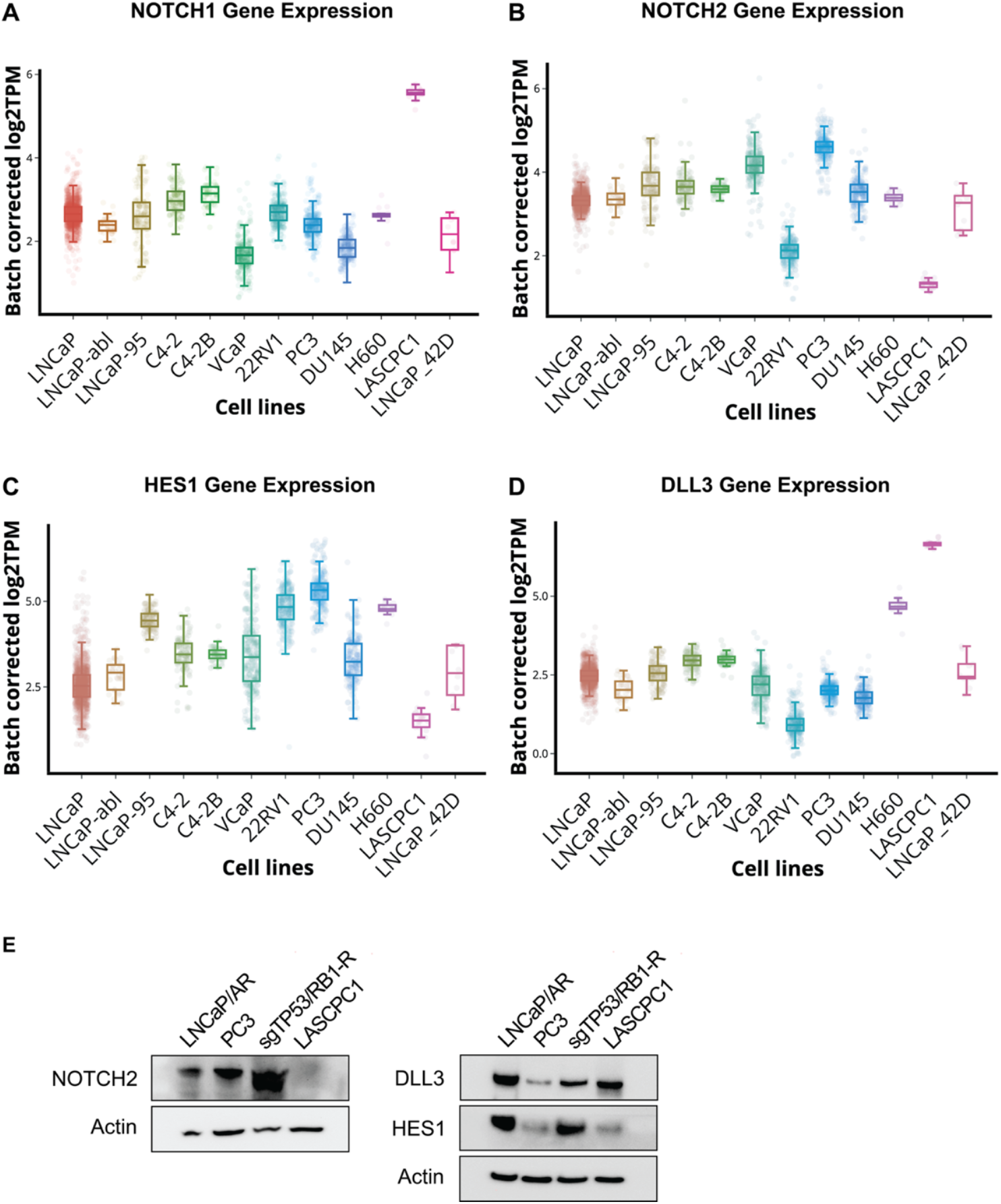
Multi-Level Validation of Notch Pathway Activation in Prostate Cancer Models. (A-D) Box and scatter plots showing bulk RNA-sequencing data for key Notch pathway genes across a panel of human prostate cancer cell lines (LNCaP, LNCaP-abl, C4-2, C4-2B, VCaP, 22RV1, PC3, DU145, H660, LASCPC1, LNCaP_42D). (A-B) NOTCH1 and NOTCH2 Gene Expression (receptor). (C) HES1 Gene Expression (downstream effector). (D) DLL3 Gene Expression (ligand). (E) Western blot analysis confirming the protein expression of NOTCH2, DLL3, and HES1 in selected cell lines (LNCaP/AR, PC3, gTP53/RB1-R, and LASCPC1), corroborating the gene expression trends observed in the bulk RNA-seq data. Actin serves as a loading control.

### TP53/RB1 Co-Deficiency Activates Notch Signaling as a Downstream Effector Driving Enzalutamide Resistance

To elucidate how Notch signaling becomes engaged during prostate cancer progression, we next investigated whether it acts as a downstream effector of TP53/RB1 co-deficiency—a hallmark genomic alteration associated with lineage plasticity and therapy resistance. While the role of TP53/RB1 loss in promoting plasticity is well established [11], its link to Notch activation has not been previously defined. We therefore examined whether TP53/RB1 loss directly induces Notch signaling. Inducible knockdown of TP53 and RB1 in prostate cancer cells (gTP53/RB1 + Dox) markedly increased the expression of *NOTCH1* and its canonical downstream target *HES1* (Figure 5A), indicating transcriptional activation of the Notch pathway upon loss of these tumor suppressors.

**Figure 5.**
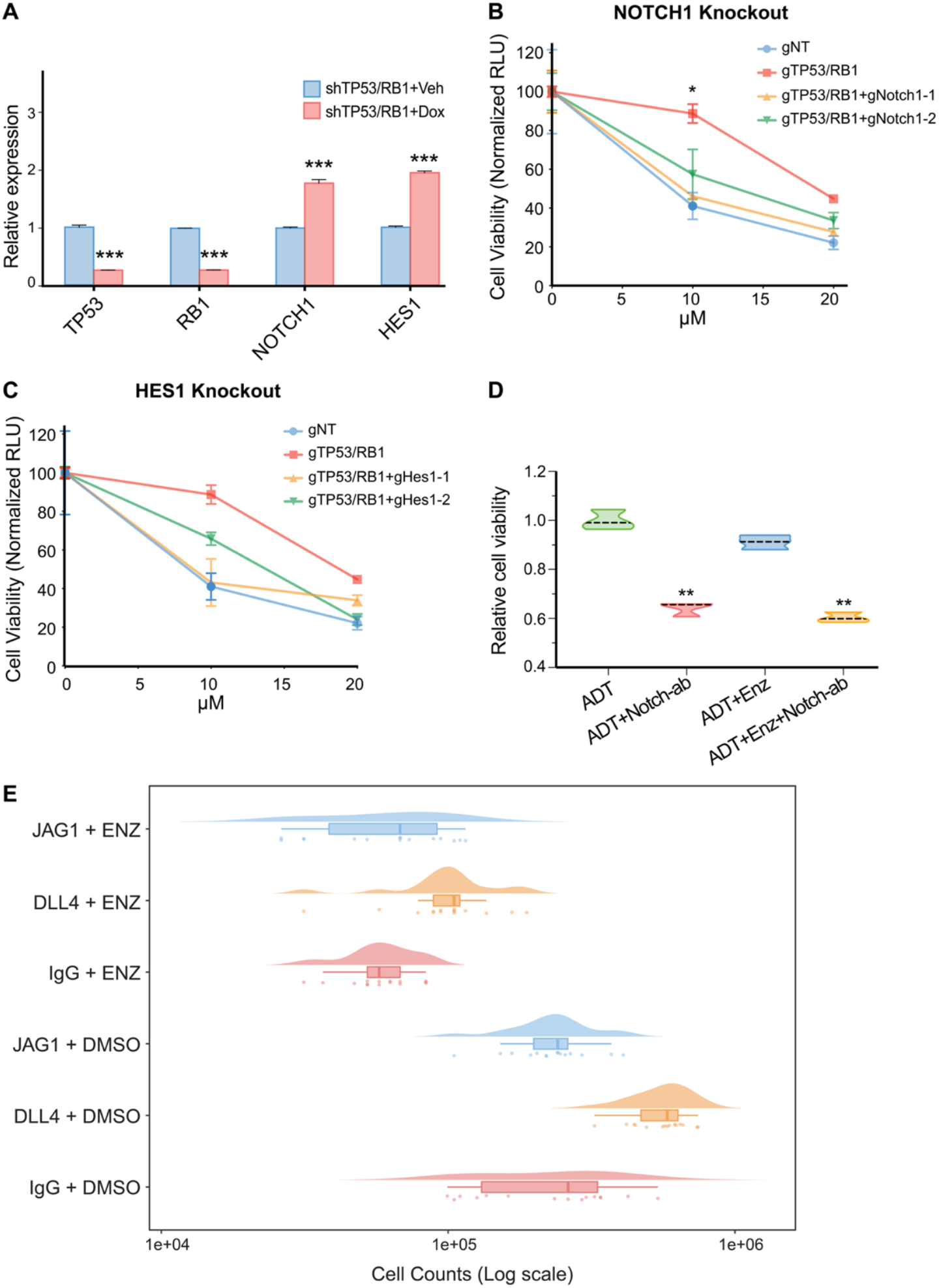
NOTCH1 and Its Downstream Effector HES1 Mediate Enzalutamide Resistance in TP53/RB1-Deficient Prostate Cancer. (A) Relative gene expression level of *TP53/RB1* downstream effectors in cells transduced with an inducible KD system. p-values were calculated by multiple t-test. For all panels, mean ± s.e.m. is represented and *** represents p<0.001. (B, C) Relative cell viability as measured by Cell Titer-Glo assay, showing cells transduced by guide RNAs targeting annotated genes. p-values were calculated by one-way ANOVA. For all panels, mean ± s.e.m. is represented and * represents p<0.05. (D) Relative cell viability as measured by Cell Titer-Glo assay, showing gTP53/RB1-ER cells treated with different combination of drugs. ADT represents androgen deprivation therapy which is achieved by CSS medium. Notch-ab represents Notch antibody. p-value were calculated by one-way ANOVA. For all panels, mean ± s.e.m. is represented and ** represents p<0.01. (E) Ligand Plate Assay. Turning on Notch signaling pathway in LNCaP/AR cells with Notch ligands (JAG1 and DLL4) show better cell viability compared to IgG control, in both DMSO control and Enzalutamide (ENZ) treated conditions. Specifically, DLL4-plated cells exhibit significantly better cell viability than IgG controls. Charcoal-stripped serum (CSS) was used to mimic ADT treatment.

To test whether this activation is functionally relevant to drug resistance, we used CRISPR–Cas9 to individually disrupt *NOTCH1* or *HES1* in the TP53/RB1-deficient background and assessed cell viability under enzalutamide treatment. As expected, TP53/RB1 double-knockout cells exhibited marked resistance to enzalutamide compared to control (gNT) cells (Figures 5B and 5C). Strikingly, deletion of *NOTCH1* or *HES1* nearly abolished this resistance, re-sensitizing cells to enzalutamide to levels comparable with control cells. These findings demonstrate that Notch signaling is a critical downstream mediator of TP53/RB1-loss–induced resistance.

Conversely, activation of Notch signaling by coating plates with recombinant Notch ligands (JAG1 or DLL4) enhanced cell viability of LNCaP/AR cells under both vehicle and enzalutamide conditions. In particular, DLL4 stimulation conferred a significant survival advantage (Figure 5E), confirming that ligand-mediated Notch activation promotes survival under androgen-deprived conditions. Together, these data establish Notch as a downstream effector of TP53/RB1 co-deficiency that drives therapy resistance through maintenance of a plastic, drug-tolerant state.

### Pharmacologic and Genetic Inhibition of Notch Reverses Enzalutamide Resistance

Although concurrent alterations in the *TP53* and *RB1* loci have been implicated in conferring lineage plasticity and therapy resistance, direct pharmacologic activation of *TP53* and *RB1* is not currently feasible [11]. This led us to investigate an alternative approach: inhibiting Notch signaling to re-sensitize prostate cancer to enzalutamide treatment in our preclinical model. While γ-secretase inhibitors (GSIs), small molecules that broadly inhibit Notch signaling, have shown preclinical promise, their clinical development faces challenges due to mechanism-based toxicities arising from non-selective γ-secretase inhibition [39]. We therefore hypothesized that a Notch1-specific antibody (Notch1-ab) could provide a more targeted and potentially safer therapeutic strategy to inhibit Notch signaling.

To create a robust model for this, we first aimed to develop a stably enzalutamide-resistant cell line from the gTP53/RB1 LNCaP/AR cells, which initially exist in a multi-lineage, plastic state with the potential to move back to AR dependency if the selection pressure is removed. These gTP53/RB1 cells were subjected to prolonged treatment with a low dosage of enzalutamide for six months to maintain selection pressure. This long-term culture under enzalutamide treatment resulted in the establishment of a derivative cell line, designated gTP53/RB1-ER (Enzalutamide Resistant). These gTP53/RB1-ER cells exhibited stable and continuous upregulation of canonical neuroendocrine lineage markers, confirming the development of a stable, therapy-resistant phenotype suitable for testing efficacy of Notch inhibitor.

We then assessed the impact of a NOTCH1 inhibitor, LEAF™ purified anti-human Notch1 antibody (Notch-ab), on the viability of these gTP53/RB1-ER cells, alone or in combination with enzalutamide, under androgen deprivation therapy (ADT) conditions (CSS medium). As measured by cell viability assay through CellTiter-Glo, the gTP53/RB1-ER cells were highly resistant to enzalutamide based on similar cell viability between ADT+Enz and ADT alone (Figure 5D). However, treatment with the Notch1 antibody significantly reduced cell viability (ADT+Notch-ab vs. ADT, p<0.01). Notably, when the Notch1 antibody was combined with enzalutamide (ADT+Enz+Notch-ab), cell viability was reduced to a similar extent as with the Notch antibody alone, and significantly more than enzalutamide treatment alone (Figure 5D). These findings demonstrate that pharmacological inhibition of Notch can overcome established enzalutamide resistance in this stably resistant, lineage-altered prostate cancer cell model, providing a strong rationale for further *in vivo* investigation.

### Notch Inhibition Drives Lineage Shift from Stem-like to Neuroendocrine-like State

We next investigated the functional consequences of Notch inhibition on lineage plasticity using both pharmacological and genetic approaches in resistant cell lines (gTP53/RB1-Resistant and PC3) (Figure 6, 7). Pharmacological inhibition using a gamma-secretase inhibitor (DAPT) or an RBPJ inhibitor (RIN1) led to the suppression of Notch pathway components (*NOTCH1/2, HES1/HEY1*). Critically, this inhibition resulted in a consistent decrease in stem-like markers (*NANOG, KLF4, JAK1*) and a simultaneous significant increase in neuroendocrine-like markers (*ASCL1, CHGA, NSE*) in both gTP53/RB1-Resistant and PC3 cells (Figure 6A, B). This suggests that Notch signaling is required to maintain the progenitor/stem-like state and that its inhibition reverses the stem-like state and promotes differentiation. This finding was validated by genetic knockdown studies using siRNA targeting *NOTCH1, NOTCH2*, or *HES1*. Genetic suppression of the Notch axis in both gTP53/RB1-Resistant and PC3 cells similarly led to a significant increase in the NE-like marker ASCL1 (Figure 7A, B). Collectively, these results confirm that suppressing Notch signaling, whether pharmacologically or genetically, alters the lineage plasticity state by pushing cells away from the stem-like/progenitor phenotype and toward a differentiated state.

**Figure 6.**
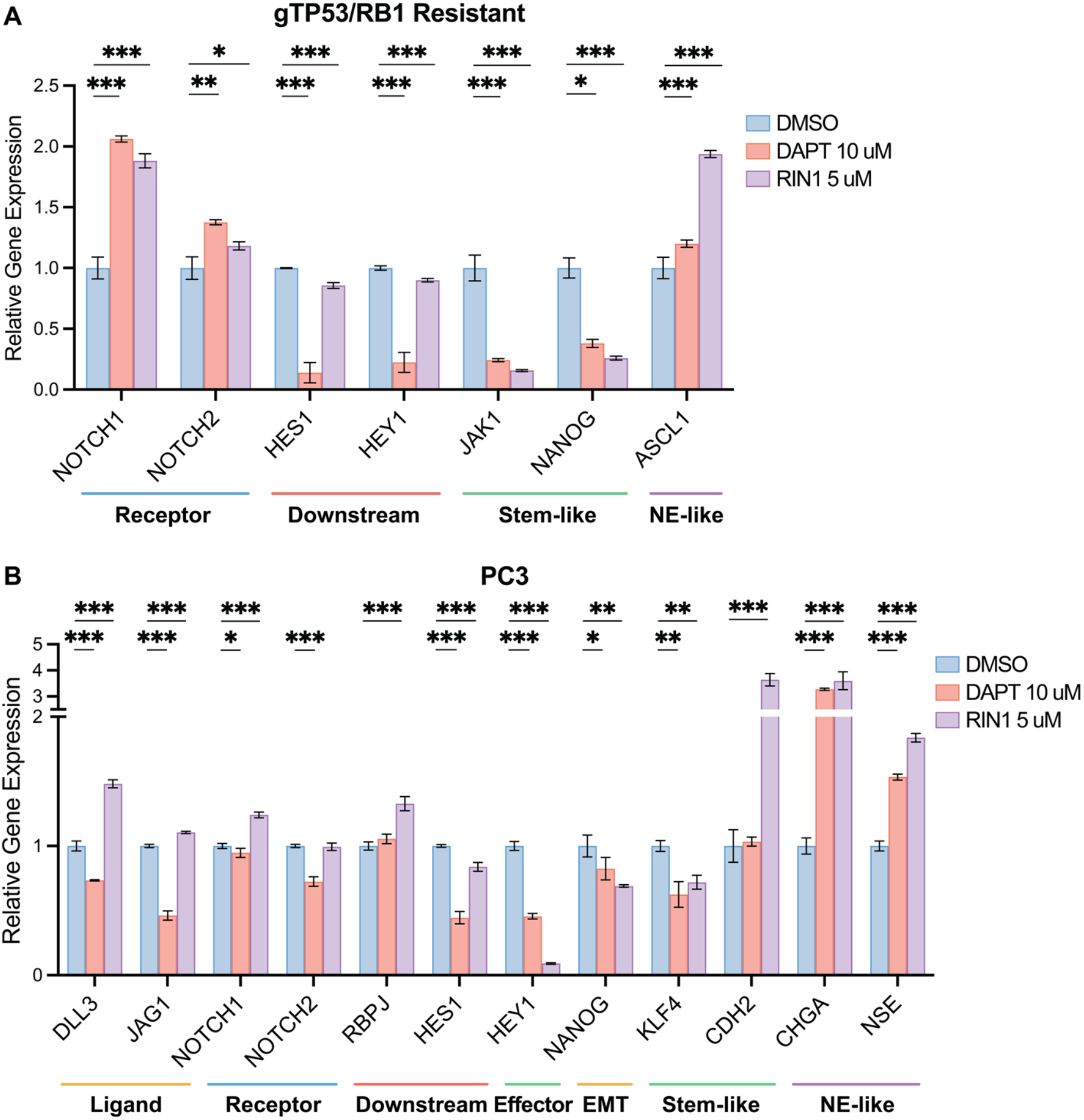
Pharmacologic Inhibition of Notch Signaling Shifts Resistant Cells from a Stem-like to a Differentiated State. (A, B) Relative gene expression of Notch pathway components and lineage markers in enzalutamide-resistant cell lines, (A) gTP53/RB1-R and (B) PC3, after treatment with a gamma-secretase inhibitor (DAPT), an RBPJ inhibitor (RIN1), or DMSO vehicle control. Significance is denoted as *p < 0.05, **p < 0.01, ***p < 0.001.

**Figure 7.**
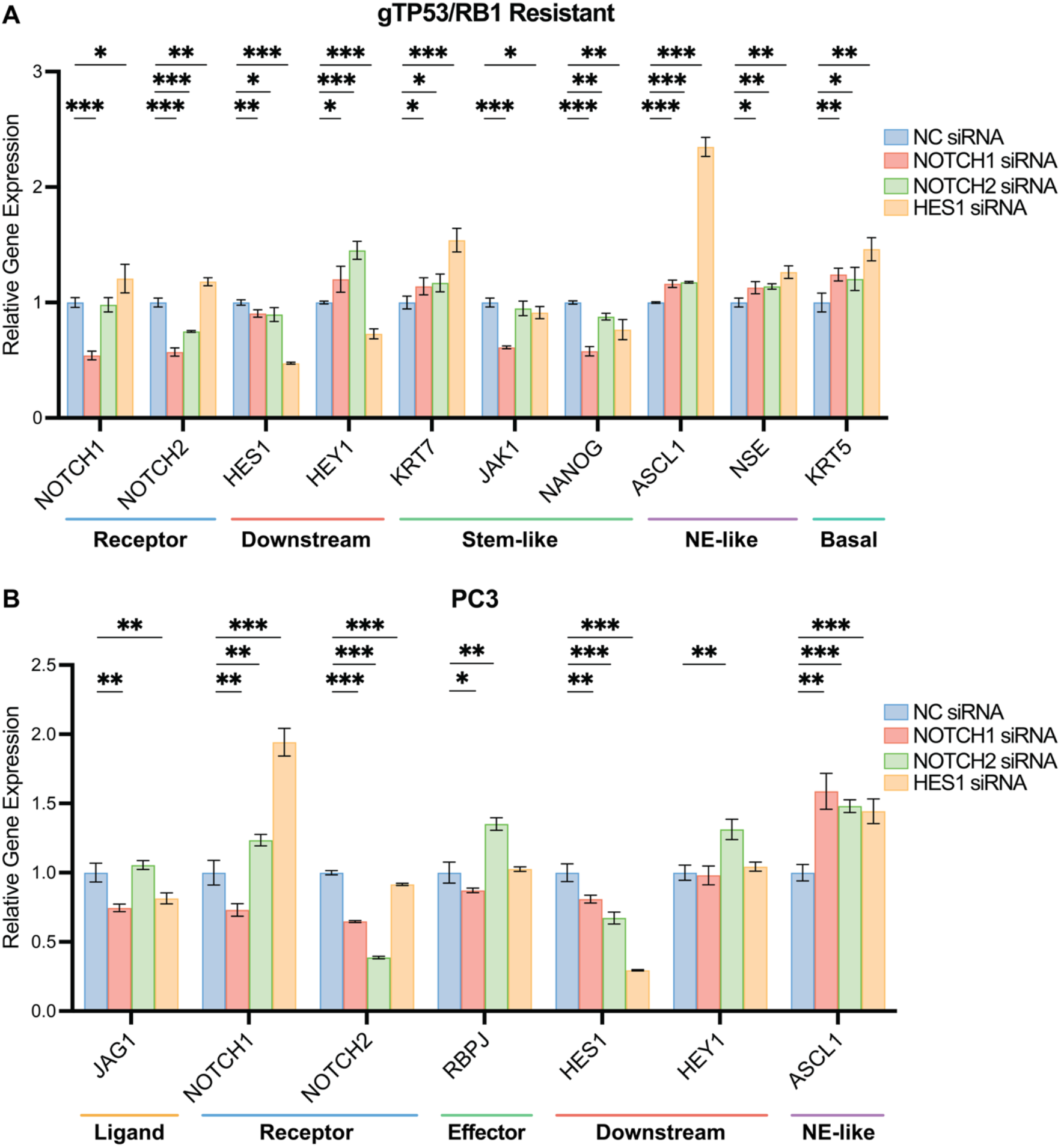
Genetic Suppression of Notch Signaling Reprograms Lineage Plasticity in Enzalutamide-Resistant Prostate Cancer. (A, B) Relative gene expression of Notch pathway components and lineage markers in enzalutamide-resistant cell lines, (A) gTP53/RB1-R and (B) PC3, after siRNA knockdown of specific genes (*NOTCH1, NOTCH2, HES1*) compared to non-targeting control (NC). Significance is denoted as *p < 0.05, **p < 0.01, ***p < 0.001.

## Discussion

The development of resistance to AR-targeted therapies remains one of the most formidable challenges in the management of advanced prostate cancer. Although next-generation AR antagonists such as enzalutamide have markedly improved outcomes for patients with CRPC, the inevitable emergence of resistance continues to drive disease progression and mortality. In this study, we uncover a critical mechanistic axis underlying this resistance by identifying Notch signaling as a key downstream effector of TP53/RB1 co-deficiency that sustains lineage plasticity and promotes enzalutamide resistance in CRPC.

Our data reveal that Notch signaling activity increases progressively during prostate cancer progression and acquisition of resistance, reaching its highest levels in therapy-resistant, AR-/NE-progenitor-like cell populations. Strikingly, this elevated activity sharply declines upon neuroendocrine transdifferentiation, suggesting that Notch signaling maintains an intermediate, metastable stem-like state that preserves plasticity but prevents full lineage commitment. Functionally, genetic disruption of NOTCH1 or its downstream effector HES1 reversed enzalutamide resistance in TP53/RB1-deficient cells, while pharmacologic inhibition of Notch using a specific blocking antibody overcame established resistance in long-term enzalutamide-resistant models. Complementary gain-of-function studies further demonstrated that ligand-mediated Notch activation enhances prostate cancer cell survival under androgen deprivation, establishing a causal relationship between Notch signaling and therapeutic resistance.

While the Notch pathway has long been recognized as a canonical regulator of stemness and differentiation, its role in prostate cancer has been controversial, with reports describing both oncogenic and tumor-suppressive functions depending on disease context. Our findings reconcile this apparent paradox by showing that Notch activity functions as a rheostat, its graded activation levels determine distinct cellular fates. Specifically, intermediate Notch activity maintains a progenitor-like equilibrium that sustains lineage plasticity and resistance, whereas excessive or diminished signaling drives differentiation toward luminal or neuroendocrine phenotypes, respectively. This concept provides a unifying framework to explain prior conflicting observations regarding Notch’s dual roles in prostate cancer biology.

Importantly, our study extends beyond correlative associations to provide direct functional evidence that Notch signaling is both necessary and sufficient to sustain therapy-resistant, stem-like states downstream of TP53/RB1 loss. Furthermore, by demonstrating that Notch inhibition, through either genetic or antibody-based approaches—restores drug sensitivity, we establish Notch as a druggable mediator of lineage plasticity in advanced prostate cancer. These results position the Notch pathway as a tractable therapeutic target that bridges genomic alterations to phenotypic adaptation and resistance.

Collectively, our findings define Notch signaling as a dynamic, tunable executor that governs the balance between stemness, lineage plasticity, and therapeutic resistance in prostate cancer (Figure 8). These insights provide a strong rationale for translational investigation of Notch inhibitors in preclinical and clinical settings. Future studies will evaluate the in vivo efficacy of selective NOTCH1 or HES1 inhibitors, explore combination strategies with AR-targeted agents, and identify biomarkers predictive of Notch dependency across diverse prostate cancer subtypes. By targeting this rheostat-like signaling network, it may be possible to reprogram therapy-resistant prostate cancer cells and restore treatment responsiveness.

**Figure 8.**
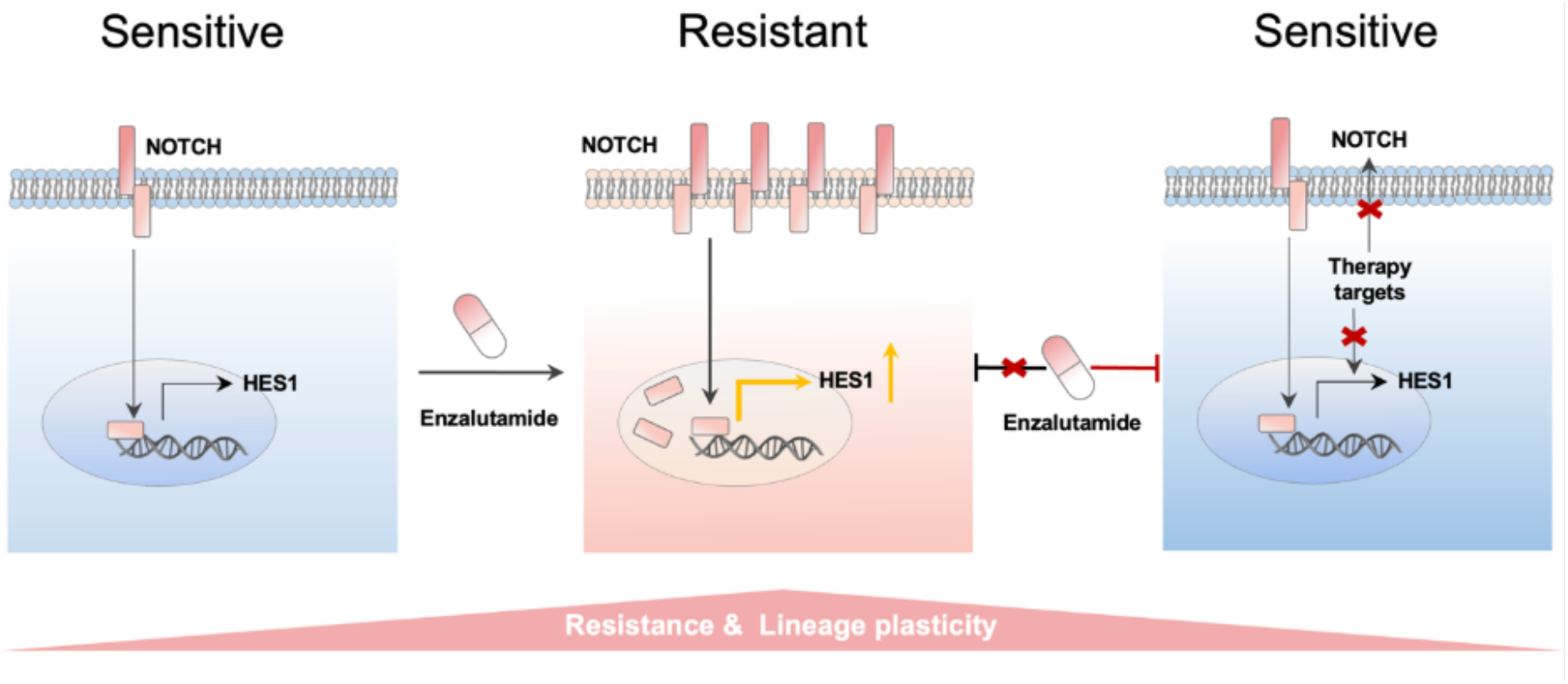
Model of Graded Notch Signaling Driving Enzalutamide Resistance and Re-sensitization in Prostate Cancer. (Left) Treatment-sensitive cells exhibit baseline Notch receptor activity and *HES1* expression. (Middle) Upon Enzalutamide treatment, resistant cells show upregulated Notch receptor expression and elevated *HES1*, associated with resistance and lineage plasticity. (Right) Co-administration of a Notch inhibitor with Enzalutamide blocks aberrant Notch-HES1 signaling, restoring treatment sensitivity.

## Methods

### Cell Lines and Culture Conditions

The LNCaP/AR prostate cancer cell line (obtained from the laboratory of C. Sawyers, MSKCC) was used as the parental line for generating experimental models of this study. LNCaP/AR cells were cultured in RPMI 1640 medium supplemented with 10% fetal bovine serum (FBS), 1% L-glutamine, 1% penicillin–streptomycin, 1% HEPES, and 1% sodium pyruvate. Cell cultures were maintained at 37°C in a humidified atmosphere containing 5% CO2 and routinely assessed for mycoplasma contamination using the MycoAlert Plus Mycoplasma Detection kit (Lonza, LT07-710), with all results being negative. Cell line identity was validated annually through human short tandem repeat profiling.

The *TP53* and *RB1* co-deficient (gTP53/RB1) LNCaP/AR cell line was generated using CRISPR-Cas9 technology as described below. For experiments involving enzalutamide treatment, gTP53/RB1 cells were cultured in RPMI 1640 medium supplemented with 10% charcoal-stripped serum (CSS medium; to maintain androgen-deprived conditions). For *TP53/RB1* knockdown induction, cells were treated with doxycycline (Dox) at 1 µg/mL for 72 hours prior to analysis. Vehicle-treated cells received an equivalent volume of Dimethyl sulfoxide (DMSO). The stably resistant gTP53/RB1-ER cell line is generated by culturing gTP53/RB1 cells in CSS medium and treating with a low dosage of enzalutamide (5 µM) for a period of 6 months.

### CRISPR and shRNA

Lentiviral-based constructs were used for CRISPR-Cas9-mediated knockout of *TP53, RB1* (for generating the initial gTP53/RB1 line), *NOTCH1*, and *HES1* in LNCaP/AR cells. LNCaP/AR cells were seeded at 400,000 cells per well in 6-well plates. The next day, medium was replaced with medium containing 50% virus, 50% fresh culture medium, and 5 μg/mL polybrene. After 24 hours, the virus-containing medium was replaced with normal culture medium. Cells were selected with 2 μg/mL puromycin for 4 days or 5 μg/mL blasticidin for 5 days. The All-In-One lentiCRISPRv2 (Addgene plasmid 52961), LentiCRISPRv2GFP (Addgene plasmid 82416), LentiCRISPRv2-mCherry (Addgene plasmid 99154), pLKO5.sgRNA.EFS.RFP (Addgene plasmid 57823), pLKO5.sgRNA.EFS.GFP, and lentiCas9-Blast (Addgene plasmid 52962) plasmids were used to generate the CRISPR and guide RNAs. Guide RNAs were designed using the Benchling guide RNA designing tool. Non-targeting guide RNA (sgNT) constructs with an empty space holder served as controls. shRNA construct LT3GEPIR (pRRL-TRE3G-GFP-miRE-PGK-PuroR-IRES-rtTA3) was originally obtained from the laboratory of J. Zuber at the Research Institute of Molecular Pathology.

### RNA Extraction and Quantitative Real-Time PCR (qRT-PCR)

Total RNA was extracted from cultured cells using Monarch® Total RNA Miniprep Kit (NEB, #T2110) according to the manufacturer’s instructions. RNA concentration and purity were assessed using a NanoDrop spectrophotometer. One microgram of total RNA was reverse transcribed into cDNA using the ABScript III RT Master Mix (Abclonal, RK20429). Quantitative real-time PCR was performed on a QuantStudio 5 System (Applied Biosystems) using the 2X Universal SYBR Green Fast qPCR Mix (ABclonal, RK21203). Gene expression levels were normalized to the housekeeping gene GAPDH. The relative expression was calculated using the comparative Ct (ΔΔCt) method. All experiments were performed with at least three biological replicates. Primer sequences are available upon request.

### Cell Viability Assay and Enzalutamide Treatment

Cell viability was assessed using the CellTiter-Glo® Luminescent Cell Viability Assay (Promega, G7570) according to the manufacturer’s protocol, with luminescence quantified on a SpectraMax iD3 automatic plate reader. All viability data were normalized to the respective vehicle-treated control cells for each cell line or condition.

#### Enzalutamide Dose-Response Curves

LNCaP/AR-derived cells, including non-targeting control (sgNT), *TP53/RB1* co-deficient (gTP53/RB1), gTP53/RB1 with *NOTCH1* knockout (gTP53/RB1+sgNotch1-1 and gTP53/RB1+sgNotch1-2), and gTP53/RB1 with *HES1* knockout (gTP53/RB1+sgHes1-1 and gTP53/RB1+sgHes1-2), were seeded at 4,000 cells per well in 96-well plates in CSS medium. After 24 hours, cells were treated with 0, 1, 5, 10, and 20 µM of enzalutamide (sourced from the Organic Synthesis Core Facility at MSKCC) or vehicle control (DMSO) for 8 days.

#### Notch Inhibitor Assays

The stably enzalutamide-resistant gTP53/RB1-ER cells were seeded at 4,000 cells per well in 96-well plates in CSS medium. After 24 hours, cells were treated with 10 µM enzalutamide, 10 µM LEAF™ purified anti-human Notch1 antibody (Biolegend, #352111), or a combination of both, as indicated in Figure 3D. Control groups received DMSO as vehicle. Treatments were maintained for 8 days.

### Ligand Plate Assay

24-well plates were coated with 25 nM recombinant human Jagged1, DLL4, or IgG1 control proteins (ABclonal; RP00877, RP00356, RPT0004) and incubated at 4°C with constant agitation for 1 hour. LNCaP/AR cells were then seeded at 10,000 cells per well and allowed to adhere at 37°C for 4– 6 hours before treatment with either DMSO (vehicle control) or 10 µM enzalutamide. After 72 hours, the culture medium (RPMI) was replaced, and the treatments were refreshed. Cells were cultured for an additional 72 hours, after which cell numbers were quantified. Each condition was performed with five biological replicates.

### Drug Treatment

0.4 million PC3 and sgTP53/RB1-ER cells were seeded in 2 mL RPMI medium in each well of 6-well plates. DMSO (vehicle control), 10 µM DAPT (MedChemExpress, HY-13027), or 5 µM RIN1 (MedChemExpress, HY-137471) were added to 3 biological replicates each. Cells were incubated under 37°C in a CO2 incubator with the inhibitors for 48 hours before being harvested for RNA extraction and subsequent qRT-PCR analysis as described above.

### siRNA-Mediated Gene Knockdown

A pool of siRNAs targeting human NOTCH1, NOTCH2, HES1, and a non-targeting control pool were used (MedChemExpress; HY-RS09445, HY-RS09448, HY-RS06134). Cells were seeded in 6-well plates and grown in antibiotic-free medium to 60-80% confluency. On the day of transfection, a 20 µM stock solution of siRNA was prepared. For each well, the siRNA pool was diluted in 100 µL of serum-free DMEM (Solution A). Separately, 6 µL of PolyJet™ In Vitro Transfection Reagent (SignaGen Laboratories) was diluted in 100 µL of serum-free DMEM and immediately added to Solution A. The mixture was incubated for 15 minutes at room temperature to allow for complex formation. The culture medium was removed from the cells and replaced with 900 µL of fresh serum-free DMEM. The 200 µL siRNA-reagent complex was then added dropwise to each well for a final concentration of 50 nM. After an overnight incubation, the medium was replaced with complete culture medium containing serum (RPMI). Cells were harvested for RNA extraction 72 hours post-transfection, and knockdown efficiency was confirmed by qRT-PCR.

### Bioinformatics

Data analysis was performed using R. Gene expression data (counts and Transcripts Per Million - TPMs) were initially loaded. The MSigDB Hallmark Notch Signaling gene set was utilized as a defined gene signature. Human gene symbols were converted to mouse orthologs where necessary using nichenetr (https://github.com/saezlab/nichenetr). Single-cell RNA sequencing (scRNA-seq) data from the Human Prostate Single-cell Atlas (HuPSA), Mouse Prostate Single-cell Atlas (MoPSA), and C42B xenograft datasets were analyzed using the Seurat V5 (https://github.com/satijalab/seurat). Standard scRNA-seq workflows included data normalization (LogNormalize), scaling, principal component analysis (PCA), and Uniform Manifold Approximation and Projection (UMAP) for dimensionality reduction. Batch correction and data integration across different studies or samples were carried out using Harmony (https://github.com/immunogenomics/harmony) as implemented within Seurat’s IntegrateLayers function. Notch pathway activity and other gene signature scores were calculated per cell using UCell (https://github.com/carmonalab/UCell). For bulk RNA-seq datasets like ProAtlas, pathway activity scores were derived using singscore (https://bioconductor.org/packages/singscore/) from TPM values. Data manipulation and wrangling were facilitated by dplyr (https://github.com/tidyverse/dplyr) and tidyr (https://github.com/tidyverse/tidyr). Visualization of results, including PCA plots, boxplots, violin plots, feature plots, dot plots, and heatmaps, was performed using ggplot2 (https://github.com/tidyverse/ggplot2), pheatmap (https://github.com/raivokolde/pheatmap), gghalves (https://github.com/erocoar/gghalves), and SCpubr (https://github.com/enblacar/SCpubr) for specialized single-cell visualizations.

### Statistical Analysis

Data are presented as mean ± s.e.m. from at least three independent experiments. Statistical significance for qPCR data was assessed using multiple t-tests. For cell viability assays, one-way ANOVA followed by Bonferroni or Benjamini correction were performed to determine significance at each drug concentration or each treatment condition. P-values for dose-response curves were calculated by non-linear regression with an extra sum-of-squares F-test. A p-value < 0.05 was considered statistically significant (*p<0.05, **p<0.01, ***p<0.001, ****p<0.0001). Statistical analyses and graphing were performed using GraphPad Prism (V9.3.1).

## Acknowledgments

This work was supported or partially supported by: National Cancer Institute/National Institutes of Health: R00CA218885, R37CA258730, R01CA288820, R01CA292949 P. Mu; Department of Defense: W81XWH-18-1-0411 and W81XWH21-1-0520 P. Mu; Cancer Prevention Research Institute (CPRIT): RR170050, RP220473, P. Mu, Prostate Cancer Foundation: 25CHAL05, 17YOUN12, P. Mu, Yale Cancer Center CCSG Pilot Grant. P30CA016359 P. Mu. and I.Y.K.

## CRediT author statement

**Yuyin Jiang**: Conceptualization, Visualization, Writing-Original draft preparation; **Siyuan Cheng**: Conceptualization, Methodology, Software, Data curation; **Catherine Yijia Zhang**: Investigation, Data Curation, Visualization; **Xiao Jin**: Formal Analysis, Visualization; **Longjun Li**: Visualization; **Melanie Fraidenburg**: Validation; **Choushi Wang**: Data Curation; **Isaac Yi Kim**: Supervision; ***Ping Mu***: Supervision, Writing-Reviewing and Editing.

## Data and code availability

The R codes necessary for reproducing all bioinformatics figures are available at https://github.com/schoo7/notch. The single-cell RNA sequencing data HuPSA and MoPSA data are downloaded from figShare: https://doi.org/10.1038/s41698-024-00667-x [39]. The bulk RNA sequencing data from patient (ProAtlas) generated by our previous publication is available through https://pcatools.shinyapps.io/HuPSA-MoPSA/ [39, 40].

